# Exploring the genetic and epigenetic underpinnings of early-onset cancers: Variant prioritization for long read whole genome sequencing from family cancer pedigrees

**DOI:** 10.1101/2024.06.27.601096

**Authors:** Melissa Kramer, Sara Goodwin, Robert Wappel, Matilde Borio, Kenneth Offit, Darren R. Feldman, Zsofia K. Stadler, W. Richard McCombie

**Affiliations:** Cold Spring Harbor Laboratory, 1 Bungtown Road, Cold Spring Harbor, NY 11724; Department of Medicine, Memorial Sloan Kettering Cancer Center, 1275 York Ave., Street, New York, NY 10065

## Abstract

Despite significant advances in our understanding of genetic cancer susceptibility, known inherited cancer predisposition syndromes explain at most 20% of early-onset cancers. As early-onset cancer prevalence continues to increase, the need to assess previously inaccessible areas of the human genome, harnessing a trio or quad family-based architecture for variant filtration, may reveal further insights into cancer susceptibility. To assess a broader spectrum of variation than can be ascertained by multi-gene panel sequencing, or even whole genome sequencing with short reads, we employed long read whole genome sequencing using an Oxford Nanopore Technology (ONT) PromethION of 3 families containing an early-onset cancer proband using a trio or quad family architecture. Analysis included 2 early-onset colorectal cancer family trios and one quad consisting of two siblings with testicular cancer, all with unaffected parents. Structural variants (SVs), epigenetic profiles and single nucleotide variants (SNVs) were determined for each individual, and a filtering strategy was employed to refine and prioritize candidate variants based on the family architecture. The family architecture enabled us to focus on inapposite variants while filtering variants shared with the unaffected parents, significantly decreasing background variation that can hamper identification of potentially disease causing differences. Candidate *d*e *novo* and compound heterozygous variants were identified in this way. Gene expression, in matched neoplastic and pre-neoplastic lesions, was assessed for one trio. Our study demonstrates the feasibility of a streamlined analysis of genomic variants from long read ONT whole genome sequencing and a way to prioritize key variants for further evaluation of pathogenicity, while revealing what may be missing from panel based analyses.

## Introduction

In the last few decades, an emerging trend across many different cancer types, both in the United States and globally, has been an increasing prevalence of early-onset cancers, generally defined as cancers diagnosed at age less than 50 (Ugai et al. 2022). While a number of potential exposures may be responsible for these trends, the impact of such exposures on genomic or genetic susceptibilities remains unknown. In colorectal cancer, a disease generally associated with advanced age (American Cancer Society, 2020), the steepest increase has been noted among individuals aged 20-29 (Siegel et al. 2023); (Bailey et al. 2015). The etiology for the increased risk of CRC in younger individuals is unknown but genetic conditions such as Lynch syndrome, Familial Adenomatous Polyposis and other rare polyposis syndromes account for only about 20% of early-onset CRCs (Cercek et al. 2021); (Al-Sukhni, Aronson, and Gallinger 2008). Testicular cancer incidence has also increased, almost doubling from 1964-2004, and it has continued to increase in the U.S. and Europe at ∼1% per year since 1992 ((Garner et al. 2005); (Nigam et al. 2015)). Testicular germ cell tumors (TGCTs) show strong heritability, with studies estimating the genetic component at nearly 50% (AlDubayan et al. 2019). Despite this, to date few genes have been strongly associated with these tumors or attributed to predisposition, with the possible exception of *CHEK2* ((AlDubayan et al. 2019); (Pyle et al. 2023)).

While the exact etiology remains unknown, it is postulated that significant changes in our exposures since the mid-20th century, such as diet, obesity, lifestyle, microbiome and the environment, may interact with genomic and genetic susceptibilities (Ugai et al. 2022). Emerging evidence suggests that the earliest phases of carcinogenesis may start early in life, with even *in utero* exposures leading to a long-lasting impact on disease susceptibility (Barker 2001); (Barker 1998). Additionally, an increase in paternal age over time, implicated as a potential risk factor for the increase in cases of autism (Wu et al. 2017); (Lyall et al. 2020); (Shelton, Tancredi, and Hertz-Picciotto 2010) has also been postulated as contributing to cancer susceptibility. In autism and other neurocognitive developmental disorders, *de novo* genetic alterations have been identified through assessment of family case-parent trios or quads (Iossifov et al. 2014); (Iossifov et al. 2012); (McCarthy et al. 2014); (Fromer et al. 2014); (Lelieveld et al. 2016). *De novo* alterations have been demonstrated in known cancer susceptibility genes (i.e., *APC, RB1, RET*, and others), and we previously have demonstrated an increased risk of de novo copy-number variants in individuals with testicular cancer (Zsofia K. Stadler et al. 2012). However, identification of structural variants in the genomes of offspring are difficult to detect with standard short read sequencing approaches and recent work from the McCombie lab and collaborators has demonstrated that long read sequencing significantly increases detection of *de novo* variants, particularly in difficult to sequence regions (Noyes et al. 2022).

Typical genomic screening techniques have relied on short read sequencing of the exome or targeted gene panels, with analysis focused on the identification of single nucleotide variants (SNVs) and small indels or large scale CNVs that can be reliably detected by such methods. However, recent papers by our group and others have emphasized the importance of structural variants in disease (Nattestad et al. 2018); (Aganezov et al. 2020); (Sudmant et al. 2015); (Abel et al. 2020). Structural variants (SVs) account for more total variation in any genome compared to SNVs (Pang et al. 2010). In addition, SVs have been linked to several diseases for which causative variants had remained elusive ((Miller et al. 2021); (Feuk et al. 2006); (Merker et al. 2018). Critically, it has been shown that long read sequencing detects many thousands of SVs per genome which would be missed by short read approaches (Nattestad et al. 2018), (Aganezov et al. 2020); (Ebert et al. 2021); (Huddleston et al. 2017); (X. Zhao et al. 2021); (Mahmoud et al. 2019). Long read sequencing allows interrogation of regions of the genome that are essentially inaccessible to short read technologies, including regions of medically relevant genes (Ebbert et al. 2019). and is now being recognized for its utility in detecting novel mutations in highly complex cancer genomes (Sakamoto, Sereewattanawoot, and Suzuki 2020). In fact, our work has shown that long reads enabled detection of structural variants in known cancer genes such as *BRCA1*, *NOTCH1* and *CHEK2* that would have gone undetected with short read methods (Aganezov et al. 2020).

Beyond SV and SNV detection, the ONT long read approach also enables the exploration of potential epigenetic mechanisms of DNA modification such as 5-methylcytosine (5-mC) signatures directly from the raw signal data, due to distinct shifts in current as modified bases transit the pore. This allows an expanded investigation that includes epigenetic regulatory factors such as promoter methylation that might impact gene expression and cancer progression. Finally, long read data allows for phasing of variants across haplotypes to enhance understanding of the larger context of multiple types of variation. This is especially useful in a study taking advantage of a family-based genetic architecture.

A striking example of the utility of long reads is shown by the latest T2T assembly of the hydatidiform mole CHM13 using long read sequencing (Nurk et al. 2022), which uncovered ∼200Mb of new sequence including 115 novel protein coding genes, as well as correcting previous reference errors which has significantly impacted variant calling accuracy (Aganezov et al. 2022). Most recently, complete phased diploid assemblies have been constructed (Porubsky et al. 2021); (Soifer et al. 2020)), and pan-genomes derived from hundreds of individuals are underway (https://humanpangenome.org/hg002/; (W.-W. Liao et al. 2023); (Gao et al. 2023). These references will provide more accurate assessment of genomic variation among individuals by resolving genotype issues that made use of previous “haploid” assemblies challenging, as well as providing a much more comprehensive representation of human diversity. Long read sequencing is significantly changing the landscape of reference genomes that have been the gold standard for many years, creating opportunities for new insights into previously unexplained disease.

As long read sequencing increases the breadth of variants discovered, it may also serve as a powerful tool in uncovering the mechanisms driving cancer predisposition in early-onset cases. Recent work has shown the importance of long reads to improve resolution of complex SVs involved in carcinogenesis (Thibodeau et al. 2020). However, methods to refine and prioritize the large number of resulting variants are necessary in order to identify events that are more likely to have functional impact.

Family studies, of which trio or quad are the simplest type, in complex diseases such as autism and psychiatric disorders (Iossifov et al. 2014); (Iossifov et al. 2012); (McCarthy et al. 2014); (Fromer et al. 2014); (Lelieveld et al. 2016) have shown the power of utilizing a family study design to filter shared variants with little impact on pathogenicity, while highlighting significant differences that may be causative. In fact, utilization of family structure has successfully uncovered drivers in pediatric cancers (Kuhlen et al. 2019); (Derpoorter et al. 2023). This may be particularly informative in early-onset cancer probands, who in addition to *de novo* mutations which have been found to be potentially disruptive and implicated in a variety of diseases (Acuna-Hidalgo, Veltman, and Hoischen 2016), may have inherited separate variants from each unaffected parent that act in concert to drive carcinogenesis.

Here, our study utilizes the power of family-based variant filtering, along with tools to select rare events which are likely to impact genomic function, to prioritize candidates for further study using extreme phenotypes of early-onset cancer cases wherein a genetic susceptibility is expected but not identified on standard targeted multi-gene assays. We demonstrate the utility of this strategy to identify a subset of interesting variants with potential functional significance, and an initial assessment of the possible impact of these variants in modulation of gene expression in matched neoplastic and pre-neoplastic lesions and normal tissue.

## Results

### Long Read Sequencing of EO-CRC Trios and Testicular Germ Cell Cancer Quad

Our study included three families with early-onset cancer proband(s). Two probands were diagnosed with early-onset CRC with germline (blood) DNA ascertained from proband and unaffected parents leading to a family-based trio design as well as tumor DNA from one of the probands. Neither of the 2 families had a family history of CRC and the probands were negative on the targeted MSK-IMPACT germline panel encompassing 90 genes associated with known cancer susceptibility (Cheng et al. 2015); (Mandelker et al. 2017). The age of diagnosis for the probands was 32 and 22 years, well below the average age of onset for colorectal cancer (American Cancer Society, 2020). The tumor of both probands was microsatellite stable, thus not suggestive of occult Lynch syndrome associated with DNA mismatch repair defects. The third family was a quad-based family structure, consisting of two brothers with early-onset testicular cancer (EO-TGCT) and their unaffected parents. In addition to also having negative MSK-IMPACT germline testing, one of the brothers had bilateral tumors at age 28, with pathological features consistent with two primary testicular cancers strongly suggestive of a genetic mechanism.

Genomes for each family member were sequenced from blood using Oxford Nanopore long reads at ∼25-35X coverage with read length N50s of ∼20-35kb per sample (Supplementary Figures 1-2 and Tables 1-8). SVs, SNVs, and methylation were then called from the aligned ONT data for all family members.

**Figure 1.**
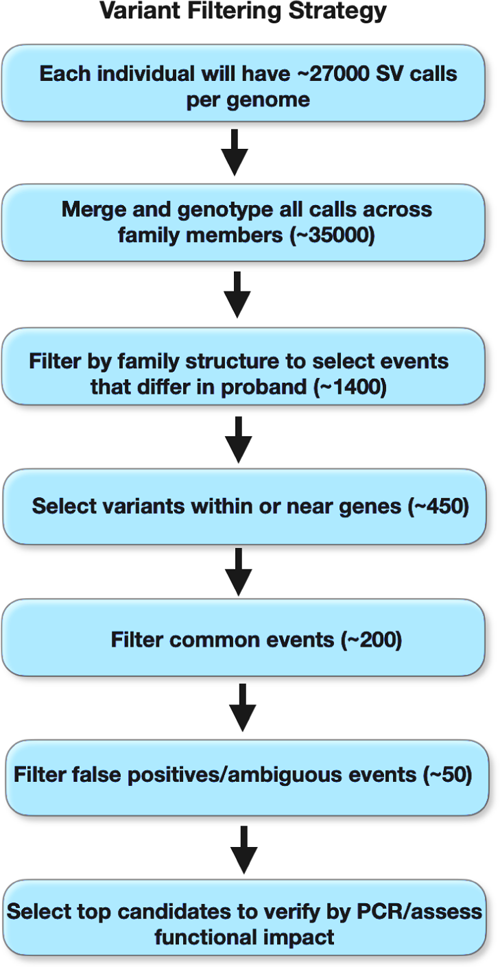
Filtering strategy to prioritize variants of interest.

### Structural variant detection

We mapped the ONT long reads to the hg38 reference using NGMLR (Sedlazeck et al. 2018) and MiniMap2 (H. Li 2018), and called SVs with Sniffles (Sedlazeck et al. 2018) to define a range of SVs including deletions, insertions, duplications, inversions and transversions. We found ∼27000-28000 SVs per individual, (Supplementary tables 3,5,8). The proportion and type of SVs within each individual was in accordance with the expected range from Chaisson et al 2019 (Chaisson et al. 2019). We then used SURVIVOR (Jeffares et al. 2017) to force genotype calls across all members of the family, in order to reduce variants that were not initially called in any individual due to the applied coverage threshold (for example, minimum total depth or number of reads showing the alternate allele). This gave us a merged set of ∼35000 genotyped variants per family (Figure 1).

SVs were then checked for Mendelian concordance with bcftools (Danecek et al. 2021), and ∼95% of the SV calls were validated. The small percentage of ambiguous calls were likely the result of sequencing error or incorrect genotyping in one of the samples. Importantly, some of these differences could be true *de novo* variants, which we explored further (see filtering by family structure section). We then used an updated method, Sniffles2 (Smolka et al. 2024), to try to improve genotyping across samples, and these SV calls had similar validation rates with a large proportion of inconsistent sites called on the sex chromosomes.

### SNP and small indel detection

It is often postulated that the higher error rate of long reads (compared to short reads) is a limitation of the technology, especially in regards to reliability of SNV and small indel detection. However, the improved mappability that long reads provide can actually improve SNV detection in complex regions (Aganezov et al. 2022). In addition, algorithmic advances have produced pipelines capable of “correcting” or reducing this raw error to allow long read detection accuracy to approach that of short reads (Shafin et al. 2021). Recent advances in the Oxford Nanopore chemistry and software have enabled improved raw read accuracy, and although this data was not produced with the latest “Q20” or duplex chemistry, additional gains would be expected with reduced modal raw error rates of ∼1%.

We initially called SNVs and small indels with the Clair pipeline (Luo et al. 2020) but found a number of false positives during manual inspection, often associated with noise in the underlying ONT reads. Increasing stringency of the Clair quality score filters to calls scored >=800 did significantly improve the results. However, we were concerned about losing true variants due to the selection of a finite threshold. Other groups have employed an ensemble approach using overlap from multiple callers to provide a more confident final variant set (Sandmann et al. 2018); (Bao et al. 2014). The PEPPER-Margin-DeepVariant (Shafin et al. 2021) pipeline uses haplotype information to improve variant accuracy. We employed this pipeline and saw improved accuracy while maintaining Ts/Tv ratios indicative of quality (∼2.04). In total we called ∼4.3 million SNVs and small indels. We see a substantial number of concordant SNV/small indel variants (85-90%) from these two callers, increasing our confidence in the variants (Supplemental Table 9). However, when breaking the calls into SNP and indel only sets, we note detection of small indels remains more difficult, as exemplified by the lower overlap of indels in each set. This is unsurprising since the error mode of ONT reads is dominated by small indels.

We then annotated the SNVs using multiple tools in Annovar (K. Wang, Li, and Hakonarson 2010) to stratify variants into coding or non-coding regions, assess population frequency in databases such as the 1000 Genomes Project (1000 Genomes Project Consortium et al. 2015) and Gnomad (Karczewski et al. 2020), predict pathogenicity of amino acid changes or splice variants, catalog conservation scores, and interrogate the variants in curated cancer databases such as ClinVar (Landrum et al. 2020), COSMIC (Tate et al. 2019) and ICGC (Zhang et al. 2019) (see methods).

### Phasing SNPs and SVs

We determined the chromosome phase of the heterozygous SNV and SV sites using LongPhase (Lin et al. 2022) which takes advantage of the long read lengths provided by ONT sequencing to create haplotype blocks, and reduces typical SNV phasing breaks by accounting for presence of SVs. The Pepper-Margin-DeepVariant variant calls and Sniffles SV calls were phased into haplotypes using the aligned long read data. We were able to achieve an average of ∼1Mb haplotype blocks when phasing the genomes individually. We did achieve some very long blocks over 20Mb as well. Haplotype blocks of ∼20Mb N50 have been reported but this is dependent upon both input read length and coverage. We then used the family pedigree info as input to WhatsHap (Garg, Martin, and Marschall 2016) to compare using the family structure to validate the proband’s predicted haplotypes in the parental genomes. Indeed when we include pedigree information we increase phasing blocks to an average of 100Mb.

### Filtering by family structure

We filtered SV and SNV calls, using the family structure, to identify variants which differed in the proband compared to the unaffected parents for each trio, or in the case of the quad, events which were shared by affected siblings but differed in the parents (Figure 1). This filtering substantially reduced the number of variants under consideration, which is especially important when assessing multiple whole genomes. Primarily, we filtered variants to those in the proximity of genes. However, importantly, we can also further assess non-coding genome regions, which may uncover larger genomic changes outside of genes. We surveyed autosomal recessive, compound heterozygous and *de novo* mutations (Supplementary Annotation Files) in the probands.

*De novo* mutations required additional QC to remove false positives in repetitive regions, and to exclude inherited events which were missed in the parents due to low coverage of either allele or mis-genotyping, particularly in the SVs. We employed the Jasmine pipeline (Kirsche et al. 2023) to refine *de novo* SV calls. Manual curation with IGV inspection of the coverage of each family member was still necessary, however this pipeline provided a reduction in false positives. We note that this careful inspection of possible de novo variants might also allow us to uncover low level mosaic variants that are present at very low read counts in one of the parents but higher levels in the proband. These potential mosaic variants could then be followed up with deeper coverage targeted sequencing or qPCR to more finely assess the presence in all individuals inclusive of potential siblings who may also harbor the identical de novo variant via parental mosaicism.

### Filtering for Overlap with Genes and for Common/Benign events

AnnotSV (Geoffroy et al. 2021) was used to assess whether SVs overlapped genes or promoters using the RefSeq annotations for hg38. Although our family structure allowed us to substantially reduce variants within each proband, we still found several hundred variants which overlapped genes. As recent studies have shown, genomic variation is widespread and common events may be found in healthy individuals with seemingly no pathogenic effect (Collins et al. 2020); (Kleinert and Kircher 2022). In order to identify events which would be more likely to be deleterious, we then checked for overlap of SVs with the dbVar (Lappalainen et al. 2013), DGV (MacDonald et al. 2014) and Gnomad (Collins et al. 2020) databases (which catalog common structural variants), to assess population frequency and prioritize rare variants. Filtering common events provided a substantial reduction in target variants. As there are many SVs yet to be cataloged, it is possible that some of our variants deemed rare might later be found to be more common, but this filtering is easily updated as databases survey more long read genomes.

### Assessing Functional Impact

Interpretation of small variants in coding regions is becoming more robust and standardized with tools to predict deleterious amino acid changes such as SIFT (Ng and Henikoff 2003), Polyphen (Adzhubei, Jordan, and Sunyaev 2013), MCAP (Jagadeesh et al. 2016) and REVEL (Ioannidis et al. 2016). In addition, one can readily assess conserved regions with tools such as GERP (Huber, Kim, and Lohmueller 2020) and CADD (Rentzsch et al. 2019), and databases of known pathogenic variants like ClinVar (Landrum et al. 2020), and COSMIC (Tate et al. 2019). SVs falling within coding regions would be likely to impact gene function and have more straightforward interpretation. However, SVs which more commonly exist outside of coding regions, have been shown to impact gene expression (Chiang et al. 2017); (Scott, Chiang, and Hall 2021). Recent studies have highlighted the importance of variation in regulatory regions and their impact on disease (Chatterjee et al. 2016); (Ignatieva and Matrosova 2021). As many of the segregating SVs we found were located within intronic regions of genes known or suspected to be implicated in cancer, we assessed overlap with ENCODE (ENCODE Project Consortium 2012) elements such as promoters, enhancers, transcription factor binding sites, DNAse hypersensitivity regions, and histone marks to increase likelihood of finding variants with functional implications, such as those that might affect gene expression. Similarly, for methylation assessment, we prioritized changes in promoter or enhancer regions, but these methods would also pinpoint methylation differences across the gene body which has been shown to affect gene expression in cancer (Q. Wang et al. 2022).

**Figure 2.**
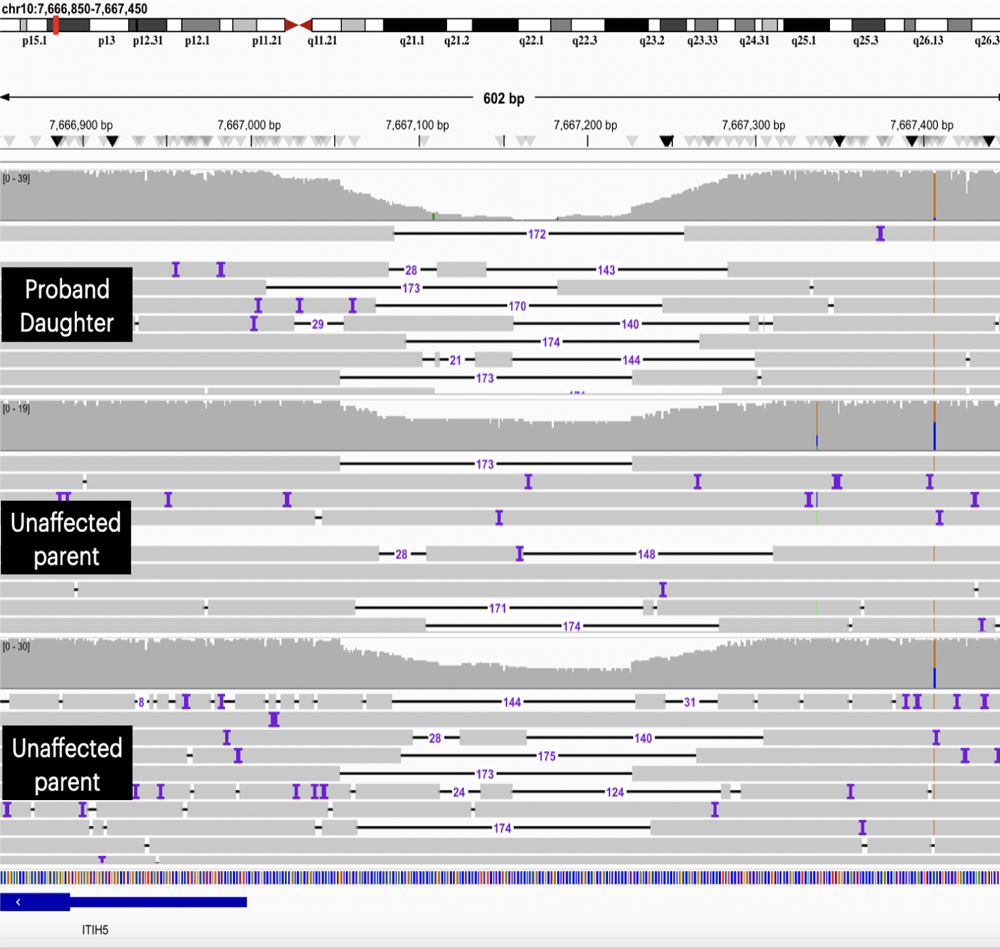
Example of a structural variant which differs in the proband compared to the parents of EO-CRC trio #2, and overlaps a regulatory region. IGV screenshot of the promoter region of the putative tumor suppressor gene ITIH5. A homozygous deletion was found in the proband (top panel), with a heterozygous deletion in both unaffected parents (middle and bottom panels).

### Methylation profiling and phasing

We called modified bases from the raw ONT signal data, specifically 5mC in CpG context, using the Nanopolish pipeline (see methods). We then used the Nanopolish calls as input to PycoMeth (Snajder et al. 2023), in order to evaluate differential methylation among the affected and unaffected samples of each family. PycoMeth assessed 1kb windows in proximity of genes, and we selected regions which showed either hyper or hypo methylation in the proband compared to the unaffected parents (Supplemental Table Z). Although we are assessing these changes in blood rather than matched normal tissue of cancer origin, we believe this may be a valuable first step which can be followed up in the appropriately matched normal tissue when available, or queried in public databases to determine if the methylation status of the gene varies across tissues.

We noted increased methylation of the promoter region of the tumor suppressor *DIRAS3* in the proband of EO-CRC trio #2 compared to the unaffected parents. Hypermethylation of promoters is often associated with silencing expression, and disruption of tumor suppressor function may contribute to cancer predisposition. We also note a region of hypomethylation in GNAS in the proband of this trio compared to the unaffected parents which overlaps an ENCODE enhancer region and may alter gene expression (**Figure 3**). Overexpression of *GNAS* has been noted in multiple cancers.

**Figure 3.**
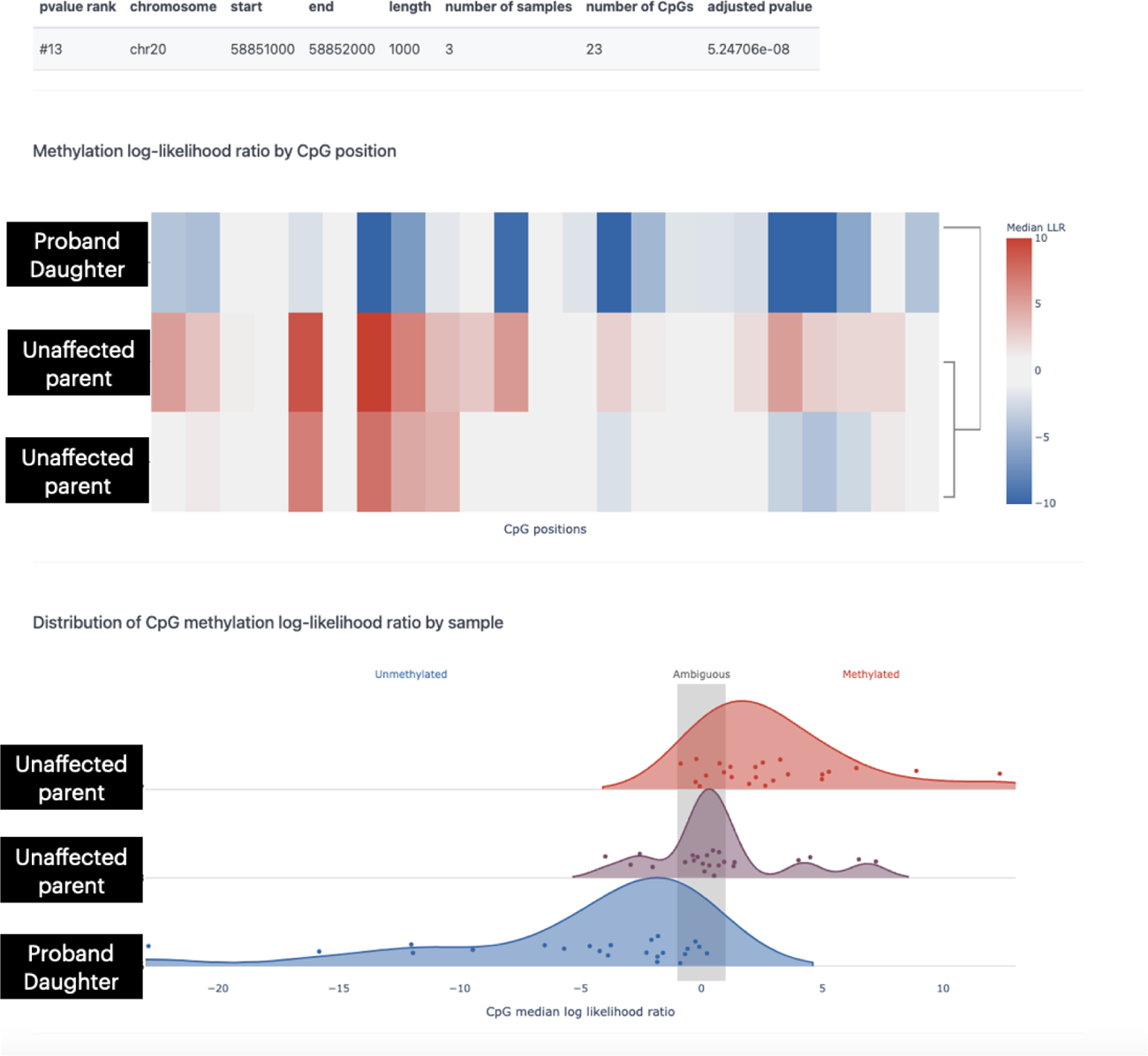
Differentially methylated promoter/enhancer region of GNAS in EO-CRC trio #2 a) PycoMeth heatmap of methylated CpG sites in windows across the 1kb region, with darker blue indicating unmethylated sites b) PycoMeth aggregated methylated CpG sites shown as dots across the 1kb region, showing decreased methylation of the proband SID103346 (peak shifted toward the left compared to parents).

In the TGCT quad, we noted increased methylation of a locus spanning an intronic enhancer in the gene *NINJ2*, and the promoter region of the overlapping antisense gene *NINJ2-AS1* (Figure 4). Overexpression of *NINJ2* has been postulated to promote colorectal cancer growth (G. Li et al. 2019), and methylation of the intronic enhancer might modulate gene expression levels. Interestingly, increased methylation in the promoter of *NINJ2-AS1* might alter expression of this long non-coding RNA. lncRNAs often modulate expression of nearby genes, such as NINJ2, but may also alter expression of other targets. In fact, predictions from the lncRNAfunc database (Yang et al. 2022) note that *NINJ2-AS1* may negatively regulate *PRMT1* expression, a gene demonstrated to be upregulated in breast, colon and other cancers (Liu et al. 2021);(Yao et al. 2021); (Y. Zhao et al. 2019). Thus, in our EO-TGCT probands, the hypermethylation of the *NINJ2-AS1* promoter may hinder down regulation of *PRMT1,* leading to cancer cell proliferation.

**Figure 4.**
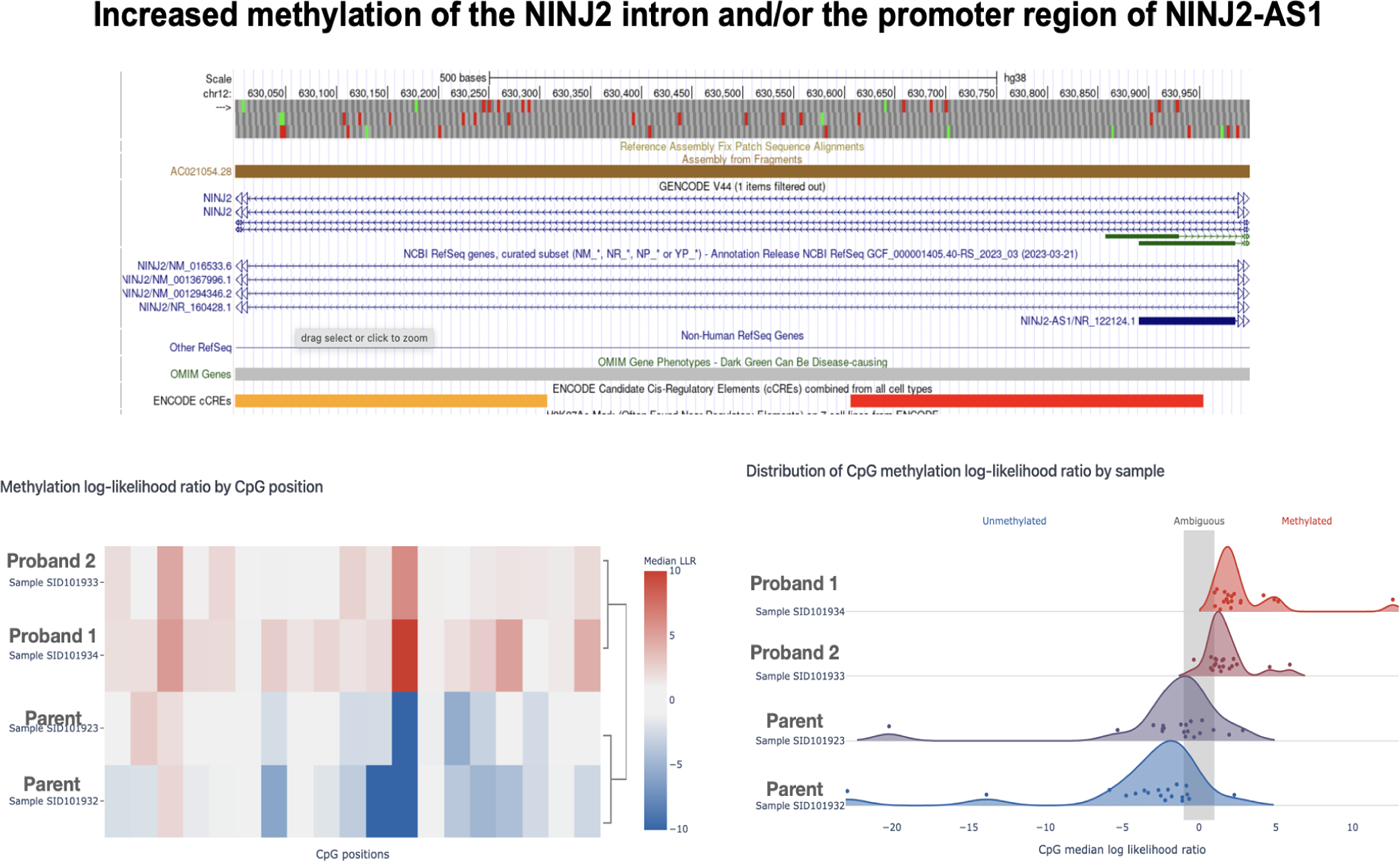
Increased methylation of regulatory elements in the NINJ2/NINJ2-AS1 region in the cancer affected sibling probands of the TGCT family. a) UCSC Genome Browser showing ENCODE elements (yellow enhancer and red promoter) in the region b) heatmap of methylated CpG sites across the 1kb region, with darker red indicating increased methylation of sites c) aggregated CpG sites shown as dots across the 1kb region, showing increased methylation of the proband (peak shifted toward the left of the grey bar indicate unmethylated CpG sites, those to the right of the grey bar indicate methylated positions).

We then used NanoMethPhase (Akbari et al. 2021) to phase the methylation and allow for allele specific analysis. The aggregated methylation calls from Nanopolish were used as input along with the phased Pepper-Margin-DeepVariant SNP calls and the aligned ONT reads. The SNP calls were phased using WhatsHap (Patterson et al. 2015) as the output format was recommended for the pipeline. We then used the DMA function to call regions where methylation was called differentially on each allele. We first noted that we could use known imprinted genes as a benchmark for assessing the ability of the tool to phase correctly. For example, we noted the *KCNQ1OT1* gene (which is exclusively expressed from the paternal allele) was detected among the haplotype specific methylation calls. Interestingly, several of the genes that were noted by PycoMeth to have differential methylation in the proband compared to the parents were found to be methylated in an allele-specific pattern. We note the possible differential methylation of the *PEG3* gene in the proband of EO-CRC trio #2, which is methylated in an allele specific manner (Figure 5). *PEG3* has been associated with the progression of colon cancer (Zhou et al. 2019); (Erfani et al. 2022).

**Figure 5.**
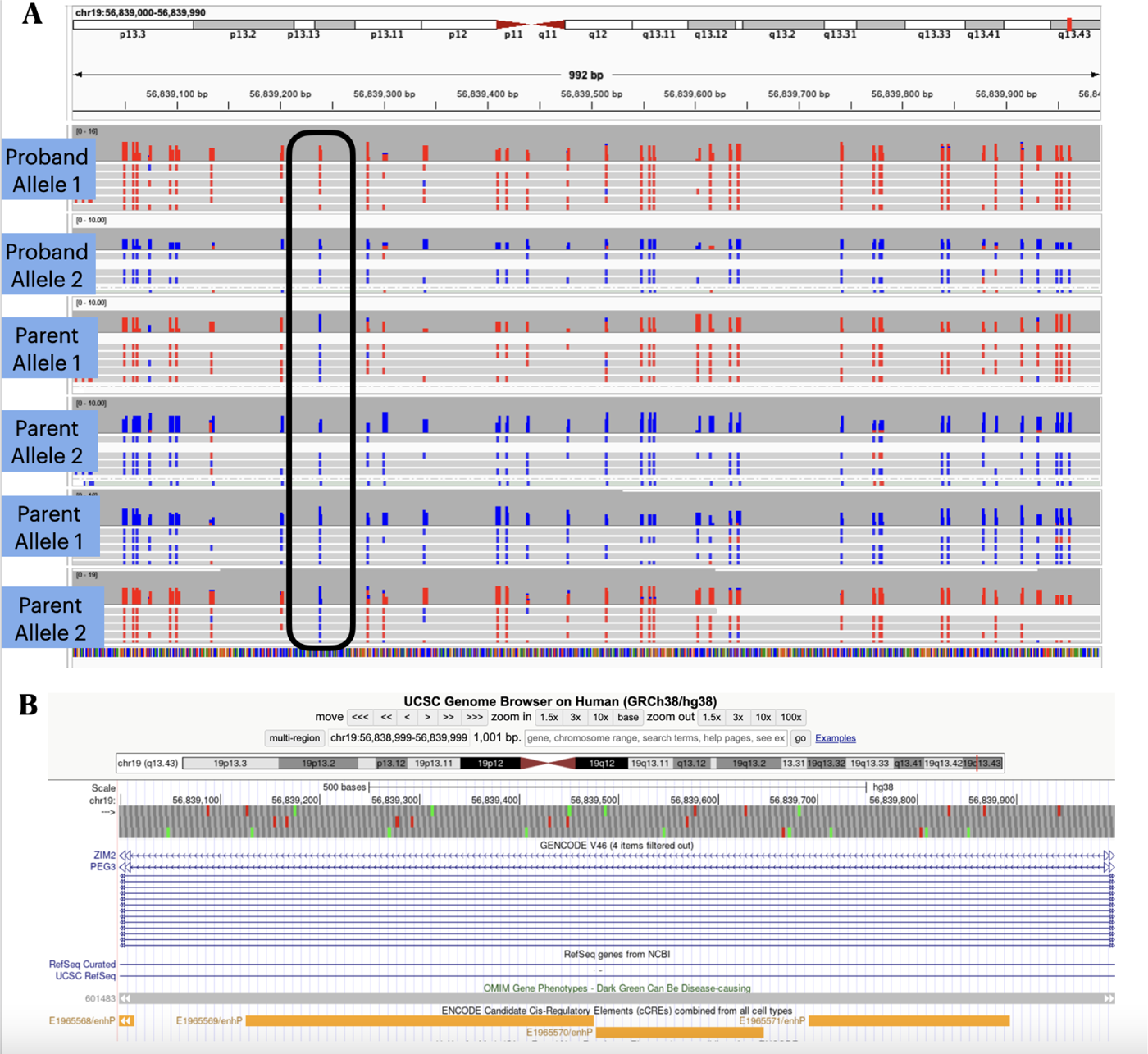
Phasing methylation provides allele specific context for PEG3 in EO-CRC trio #2. a) IGV screenshot showing phased alleles of the proband and parents. In “bisulfite mode”, red colored bases represent methylated Cs and blue indicates unmethylated Cs. Each family member contains one methylated (silenced) allele in red, and one expressed allele. Black encircled CpG site is unmethylated in all parental alleles but methylated in one proband allele. b) UCSC Genome browser showing overlap with ENCODE enhancer marks.

### Prioritized Variants

We have uncovered multiple variants of interest in the probands (Table 1). The EO-TGCT probands were found to have a homozygous deletion in the last exon of the gene *ZNF738* (Figure 6) which contains a KRAB domain and may be involved in transcriptional regulation. KRAB Zinc finger proteins have been shown to regulate many key cancer pathways such as p53 and Wnt (Sun et al. 2022). We include these autosomal recessive cases since variants of this type might exhibit a range of functional impact depending upon penetrance and epistatic interactions with other genomic alterations. Indeed, over the last decade with the advent of next-generation sequencing, a number of autosomal recessive cancer predisposition syndromes have been discovered where the heterozygous state confers no or minimal cancer risk, but the homozygous state is associated with a high-penetrance cancer syndrome (*MUTYH, NTHL1, MSH3, MLH3, BLM, ERCC1/2/3, FANCA/C/G* among others*) (Garutti et al. 2023)*.

**Figure 6.**
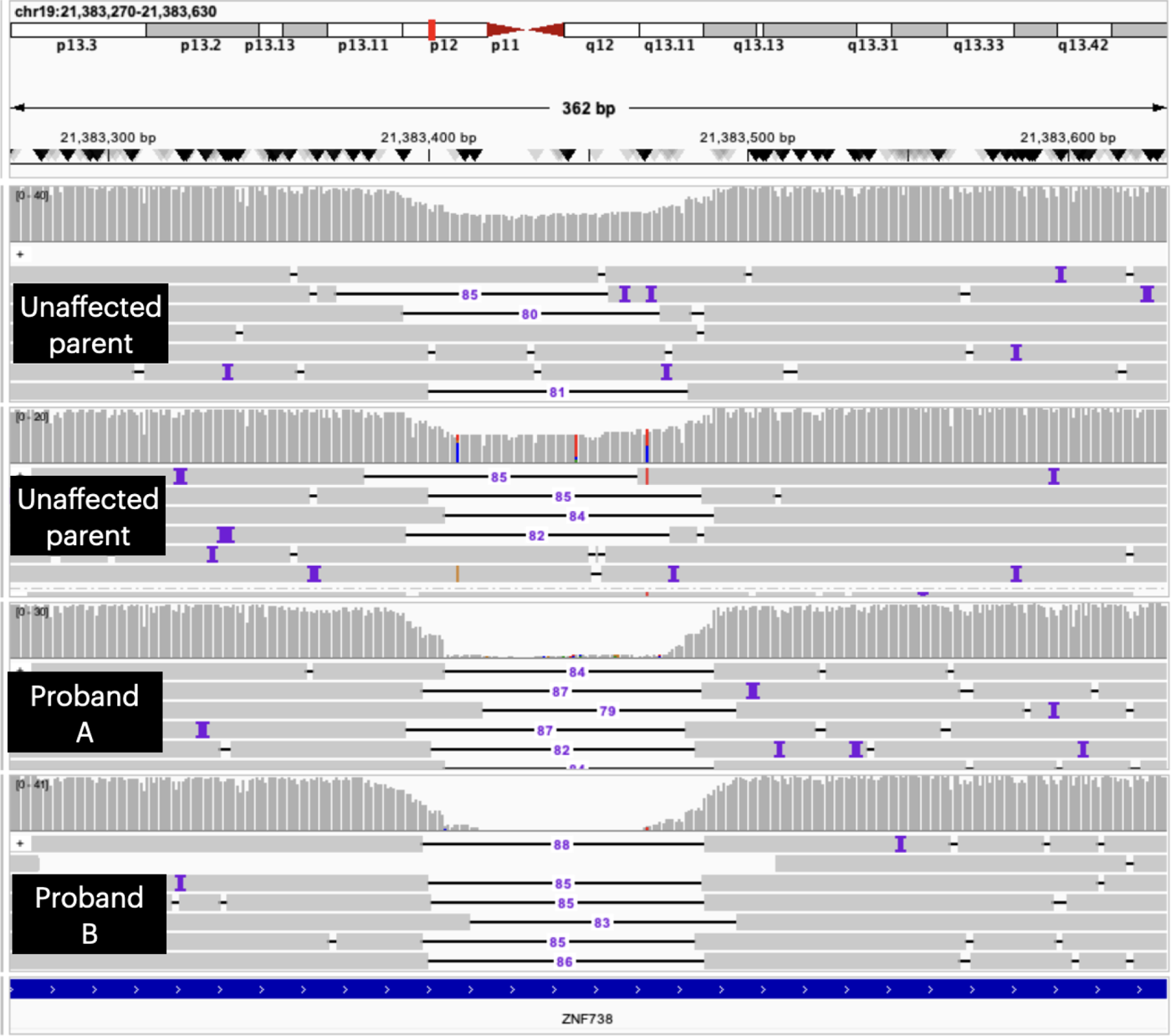
IGV screenshot of the last exon of gene ZNF738 in the TGCT quad. Aligned reads are displayed with heterozygous deletions in the unaffected parents (top panels), but homozygous deletions in both probands (bottom panels).

In the EO-CRC trios, we noted SVs in genes related to cancer, such as *TP73* and *RASA3*, and in genes implicated in CRC in particular (such as *ITIH5, SMYD3, BCL2*). Several of these variants overlap with regulatory elements and may change expression of these genes. For example, a compound heterozygous deletion in EO-CRC trio #1 in *BCL2* overlaps an enhancer region. Rearrangement of this enhancer might lead to altered expression of this gene, whose overexpression and deregulation have been linked to prolonged survival of cancer cells (Qian et al. 2022) and CRC progression (Ramesh and Medema 2020). Another example is a compound heterozygous mutation in EO-CRC trio #1, which abuts an ENCODE annotated enhancer region (ENCODE Project Consortium 2012)) in *TP73* (Figure 7). TP73 is part of the p53 family of tumor suppressors and is well-known to be downregulated in CRC (Kotulak et al. 2016). Investigation of functional consequences of this large class of until recently relatively poorly characterized variants may uncover mechanisms of “missing heritability”.

**Figure 7.**
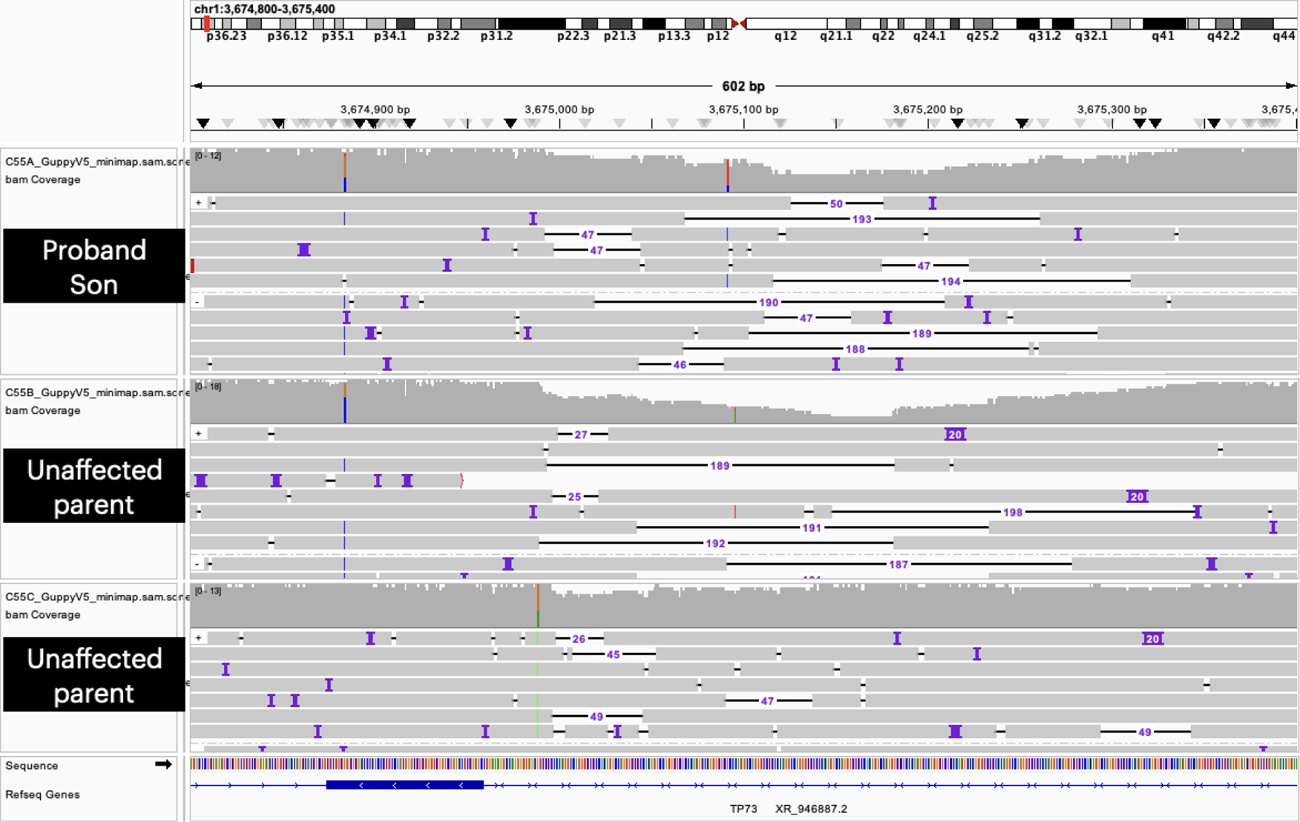
Compound heterozygous SV in EO-CRC trio#1 in the gene TP73. IGV screenshot of aligned reads with a heterozygous larger deletion in the father (middle panel), a heterozygous smaller deletion in the mother (bottom panel) and both deletions in the proband (top panel).

We have also identified differential methylation of promoter regions in genes such as *DIRAS3* (a putative tumor suppressor) and *GNAS* (an oncogene), which may alter expression and contribute to the progression of the disease in the proband. We also note that although the hypermethylation of the additional *GNAS* enhancer region is not the canonical model for increasing oncogenic expression, recent studies have shown that hypermethylation may in fact have activating effects as well as the more typical suppression (Smith et al. 2020). Many of these changes can have varying effects, and their consequences are only beginning to be explored.

To address the potential effect of these events, we performed expression analysis of blood and colon tissues and tumor which were available from EO-CRC trio # 2 (see section below).

**Table 1.**
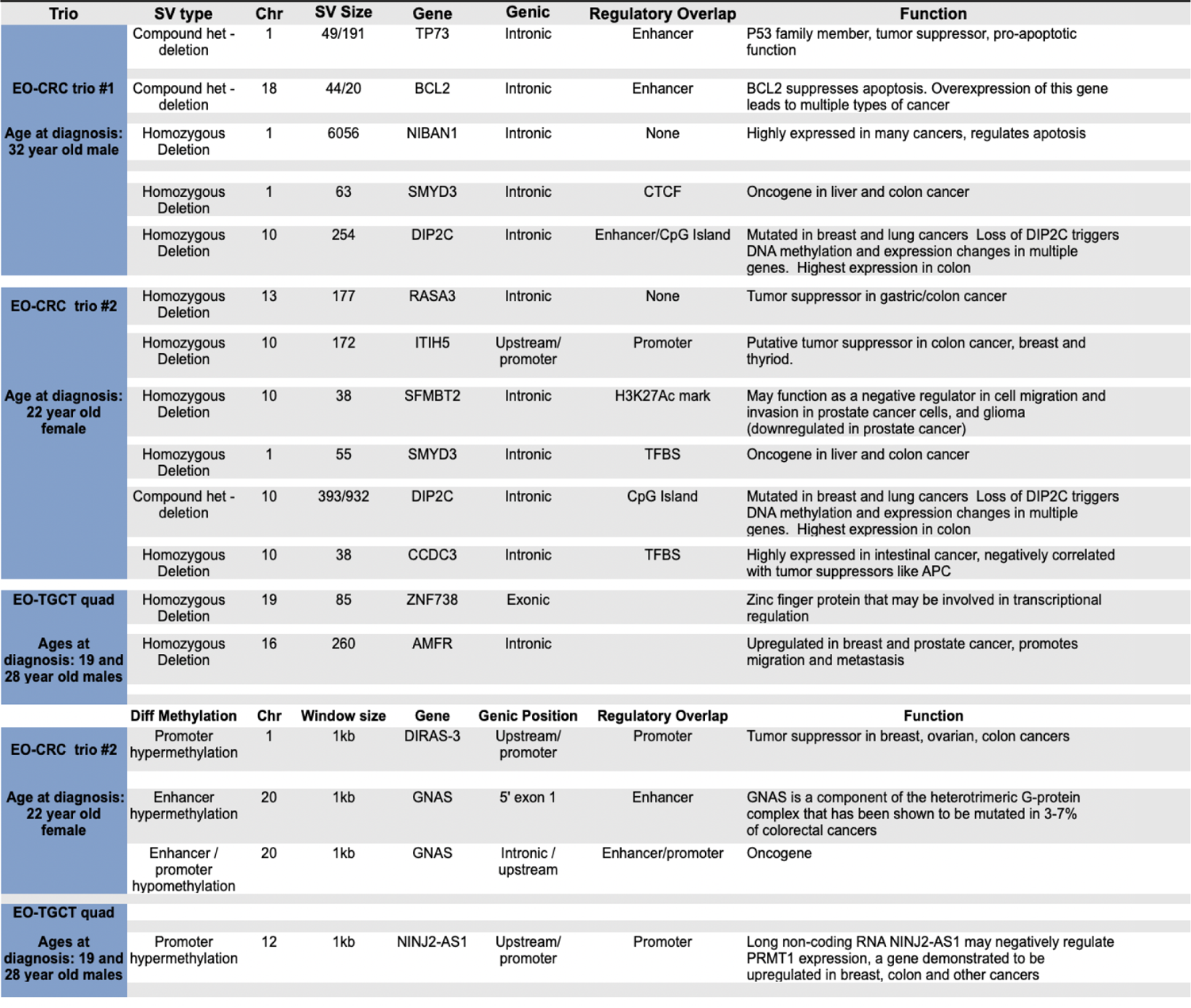
Selected structural variants and methylation changes found in each family that may be involved in cancer predisposition. Overlap with regulatory elements and possible function of the genes in cancer are noted.

### Additional information from the long read CHM13 genome

In order to utilize the more complete reference genomes now available, we mapped our long read data to the CHM13 reference. We were able to identify variants that fell in regions that were corrected in or added to CHM13 compared to hg38. In some cases, these variants would have been missed by utilizing hg38. A homozygous deletion in the *RPH3AL* gene in EO-CRC trio #1 (Supplementary Figure 3) falls in a region of CHM13 that is unique compared to hg38, with whole genome alignments using the Cactus tool (Armstrong et al. 2020) showing the altered regions between the references. This is a likely tumor suppressor gene downregulated in bladder cancer (Ho et al. 2012). A homozygous deletion in the *ZNF793* gene in EO-CRC trio #2 falls in a region of CHM13 where Cactus alignments show a break in alignment to hg38 (Figure 8). This partially overlaps an exon of this gene which has been associated with gastric (Gómez et al. 2021) and esophageal cancers (Yu et al. 2015). These examples highlight the importance of long read references for resolving regions of genes that might be functionally relevant.

**Figure 8.**
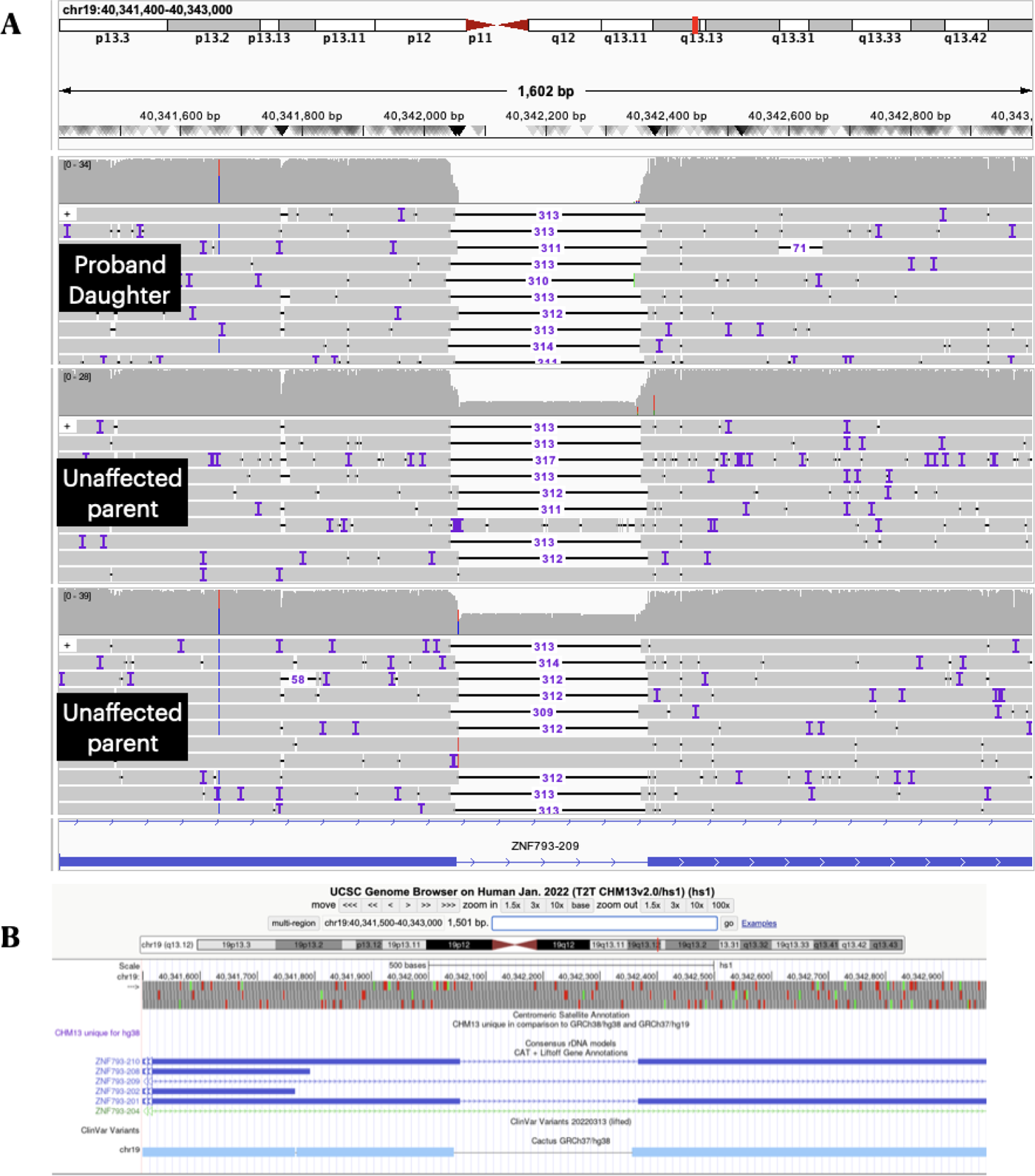
Homozygous deletion overlapping an exon of ZNF793 in EO-CRC trio #2. a) IGV screenshot of T2T-CHM13 aligned reads with a homozygous deletion in the proband, and heterozygous deletion in both unaffected parents. b) UCSC Genome Browser view of the ZNF793 exon region in CHM13 with bottom track showing break in alignment of CHM13 compared to hg38.

### Profiling expression in tumor, matched normal adjacent and adenomatous polyps

In order to ascertain possible transcriptional effects of the prioritized SVs we performed Illumina short read RNAseq on one of the EO-CRC probands to quantify gene expression. Germline SVs have been shown to correlate with transcriptional changes in tumors from multiple tissue types (Chen et al. 2024).

We prepared 3 replicates of FFPE sections for the proband of EO-CRC trio #2 including colon tumor, normal adjacent colon, and adenomatous polyps (Figure 9).

**Figure 9.**
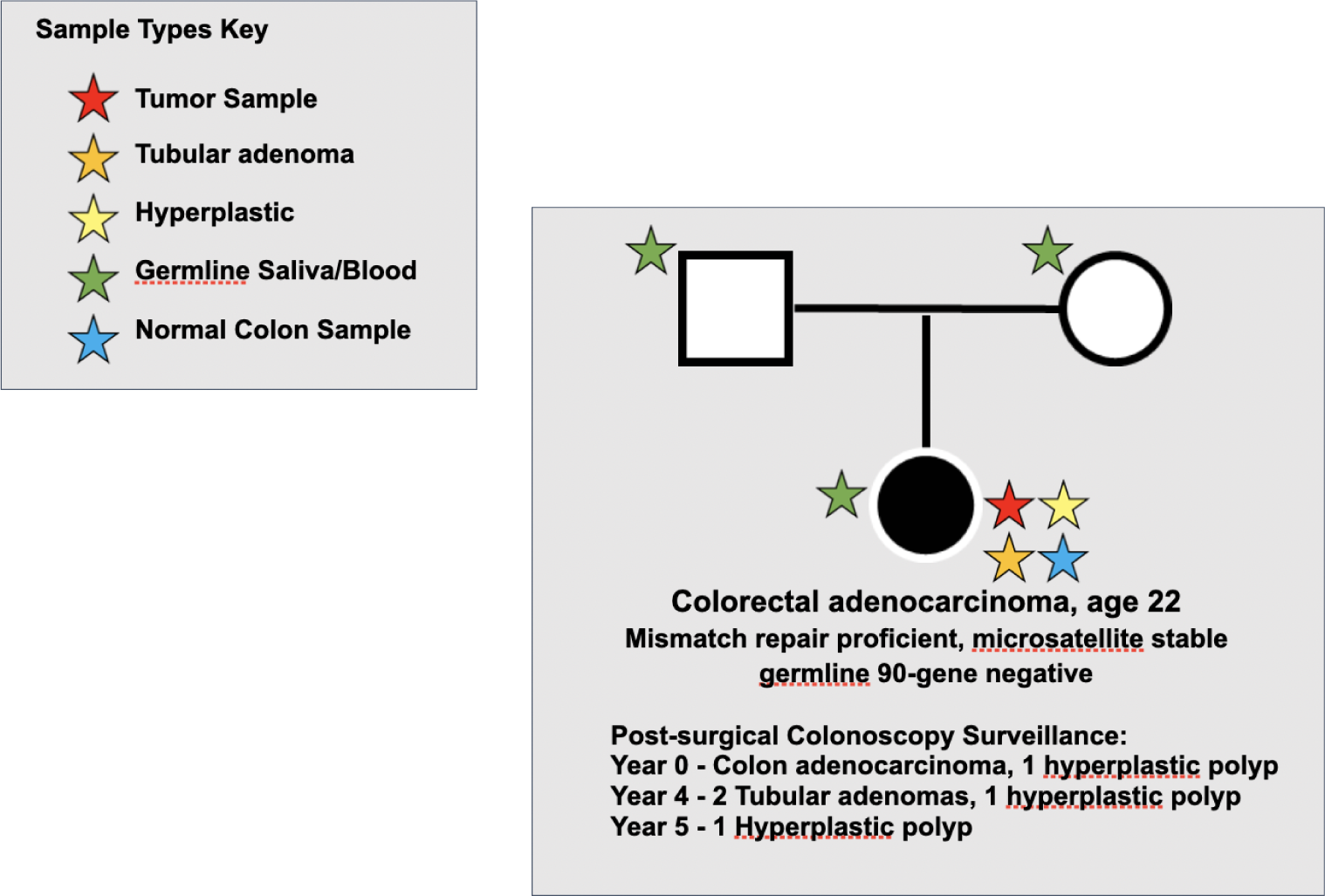
Pedigree of EO-CRC #2 noting multiple samples ascertained for the proband. For the 22 year old proband we performed germline (blood) long read whole genome sequencing, as well as RNAseq from FFPE sections of normal colon, 2 adenomatous polyps, and the colon tumor.

We aligned these reads to hg38 and normalized read counts across genes for each sample using DESeq2 median of ratios (Love, Huber, and Anders 2014). Availability of such normal and pre-neoplastic colonic tissue sampling provides several advantages. For several putative functional variants identified, we are able to demonstrate the change in expression in the proband from normal colon to the adenomas, and between normal colon and the tumor (Supplemental Figure 4). For example, *DIP2C* has reduced expression in breast cancer (J. Li et al. 2017), and we found a compound heterozygous SV in this gene. Expression is reduced from the normal adjacent colon to a lower level in the adenomas, and expression is also reduced in the adenocarcinoma compared to normal colon. We found a homozygous deletion in the proband which overlaps the last exon of *ZNF793*, a potential tumor suppressor (Yu et al. 2015), and also shows progressively decreasing expression from normal colon to adenoma to adenocarcinoma. Similarly *ITIH5*, a tumor suppressor inactivated in CRC, (Kloten et al. 2014), and in which the proband harbors a homozygous deletion in the promoter region, decreased expression is demonstrated in the adenomas and tumor compared to the matched normal. It is important to point out that although we profile the expression of specific genes containing variants in these examples, many of these alterations may also impact the expression of other genes in up or downstream networks implicated in cancer. Indeed, loss of *DIP2C* has been shown to impact expression of the tumor suppressor *CDKN2A* among many other genes (Larsson et al. 2017). As such, exploring variants in genes not typically associated with cancer may uncover a new contribution to carcinogenesis through regulatory interaction with genes or pathways previously implicated in cancer.

We recognize that factors such as developmental stage and oncological treatment may also affect expression. Notably, the EO-CRC proband’s adenocarcinoma assessed in this analysis was obtained prior to the patient receiving any chemotherapy or radiation treatment. The pre-neoplastic adenomas developed >3 years after completion of oncological treatment and were identified during standard colonoscopic surveillance. We acknowledge that this is a first tier assessment of the potential functional impact of these variants and additional validations would be expected downstream. While acquired genomic changes in the adenocarcinoma may also further contribute to differences in expression, our analysis does not preclude the possible importance of germline variants in the progression of these changes. Moreover, our analysis of not just germline but also pre-neoplastic and neoplastic lesions from the same individuals serves as a powerful tool enabling further prioritization of variants.

## Discussion

Although our understanding of cancer susceptibility has significantly improved with the use of next-generation sequencing technologies, to date, most clinical and even gene discovery efforts have focused on the the utilization of multi-gene panels or exome-based sequencing using short read sequencing focused on detection of point mutations in the coding regions of genes. These explorations have led to ∼20% of early-onset cancers being explained by genetic susceptibility, a significant enrichment over average onset cancers (Mandelker et al. 2017) (Zsofia Kinga Stadler et al. 2023). Nonetheless, the vast majority of early-onset cancer patients have no identifiable genetic risk marker. A lack of understanding of cancer etiology in these patients results in ambiguity with respect to preventative cancer surveillance not just in the cancer survivor but also in potentially at-risk family members. It may well be the case that the major rare and higher-penetrance cancer susceptibility genes have already been identified. However, the variants which alter these genes, or regulate the expression of these genes, may only have been superficially sampled and the role of epigenetic and other protein changes that modulate their expression are yet to be elucidated.

Additionally, although small nucleotide changes have been well characterized, thousands of SVs and their effects, particularly on gene regulation, have yet to be explored. Similarly, canonical methylation patterns (for example, promoter hypermethylation for gene silencing) have been well studied, but a deeper understanding of the full range of epigenetic changes in known cancer genes that do not harbor typical disruptive point mutations may yield new insights into epigenetic regulation of gene expression. Importantly, the larger view of variation across haplotypes may uncover “lower impact” variants that act together to synergize effects, much as distant SVs may cause downstream methylation changes that alter expression. Assessment of the full spectrum of germline variation is needed to more fully understand the role of these alterations in the genome and their possible contribution to cancer susceptibility.

Long read sequencing may unlock a host of previously undetected variation at both the genetic and epigenetic levels. Mining this rich resource may provide clues to the mechanisms driving currently unexplained cancer predisposition. A framework to filter and rank the many variants of unknown function that will be discovered using family based studies may streamline prediction of candidates most likely to impact disease.

Our study shows that a family-based filtering strategy enables identification of regions of significant genomic difference between the affected individuals and their unaffected parents, providing a computational sieve to catalog variants of interest which have been less well studied, and which may shed light on additional mechanisms that may perturb known cancer pathways. This filtering strategy reduces the number of structural variants under consideration by approximately 100 fold (from ∼30,000 shared family SVs to ∼300 differing SVs), and highlights several regions of differential methylation with proximity to genes. A gene list of this size then becomes more manageable to study in a tiered fashion. Importantly, our approach of analyzing not just germline lymphocyte-derived DNA but also normal tissue from the at-risk organ (i.e., normal colonic tissue), in addition to preneoplastic and neoplastic lesions from the EO-CRC trio demonstrates a powerful and unique technique for further prioritization of putative functional variants through assessment of altered gene expression. Downstream functional analyses, often requiring significant resources, may then focus on the most promising candidate variants.

Since the advent of exome sequencing (Hodges et al. 2007), which was originally envisioned to reduce the cost of sequencing to a manageable level, the cost of whole genome sequencing has dropped more than a hundred fold. Even whole genome sequencing with long read instruments has dropped 50-100 fold since their development. Rather than focusing solely on exonic regions, a whole genome survey provides opportunities to integrate both coding and non-coding variants, and to investigate the interplay of variants which may elucidate novel mechanisms of cancer susceptibility. However, functional analyses of these candidates such as CRISPR screens in cell lines or organoid/mouse models require substantial time and effort, highlighting the importance of establishing a candidate variant list with the highest potential for pathogenicity. Though no tool can perfectly predict which genomic variants may contribute to disease, an approach such as ours that utilizes a family-based architecture, assesses multiple types of genomic variation, and then carefully prioritizes likely functional alterations promises to yield the most fruitful and efficient approach for downstream functional analysis. We also demonstrate the flexibility and utility of our pipeline on both different cancer types and in different family structures. These investigations, and more like them, at a larger scale, can improve our understanding of the variety of changes that may alter genomic function and predispose individuals to carcinogenesis. At this point, the analyses of data like this are rate limiting, but that challenge can be addressed by larger studies with more functional analyses.

## Materials and Methods

### Patient Samples

Germline DNA from blood was obtained from two case-parent trios and one case-parent quad under research protocols approved by the Institutional Review Board of MSK with written informed consent obtained from each study subject. In the cancer affected probands, if possible, other tissue, inclusive of pre-neoplasia and neoplasia was also ascertained. Selection of case-parent trios/quads included individuals with early-onset cancers without an explained genetic etiology, or as in the case of the TGCT quad, the presence of siblings with the same rare cancer type suggestive of an underlying genetic predisposition.

### Sample preparation and ONT sequencing

Blood (∼10mL) collected at MSK was shipped to CSHL and stored at 4C to preserve DNA integrity. DNA was extracted with the Circulomics ultra long kit. DNA was assessed for quality via Qubit, Nanodrop and Femto pulse, to measure mass, purity and size, respectively. DNA was prepared for sequencing with an ONT SQK-LSK109 kit. The libraries were sequenced on a PROM0002 PromethION cell. Approximately ∼20-40x sequence coverage from each individual (Supplementary Tables).

### ONT base calling and data processing

ONT data was base-called in real time during the sequencing run using the ONT GuppyV4 basecaller with the high accuracy model. Data was then re-processed to improve quality using GuppyV5 with the “super” high accuracy model. Run metrics were tracked for each flowcell, including total yield, percentage of data passing quality filters, average/N50 read lengths, and proportion of data in reads >30kb. This ensured appropriate quality and targeted coverage levels for each sample. ONT fastq files were transferred to local storage on the CSHL HPCC for further processing. Larger raw signal data (fast5 files) were archived to storage arrays and transferred to the active storage as needed.

### ONT sequence alignment and reference comparison

The long read fastq files were aligned to the hg38 human reference genome using the NGMLR(Sedlazeck et al. 2018) aligner with parameters tuned for long read ONT data and with Minimap2 (H. Li 2021). For the long read CHM13 reference, reads were aligned using NGMLR to the CHM13v1.1 release. BAM files were sorted with Samtools and aligned depth was calculated.

### Structural variant detection

SV calls were ascertained with Sniffles(Sedlazeck et al. 2018), using a minimum read support for the alternate allele of 6. Because SV breakpoints may often be shifted for the same event in different samples, SURVIVOR (Jeffares et al. 2017) was used to merge SVs within a 1kb window and create a merged genotyped variant file for each family. Variants were then filtered by the family structure. Jasmine was used to increase accuracy of predicted de novo SVs and Sniffles2 was run to improve genotyping.

### SNP calling

SNVs and small indels were called with Clair (Luo et al. 2020), and filtered for quality score >800 in order to remove low confidence calls. SNVs and indels were then filtered according to the family structure as described above. SNVs were then re-called with the PEPPER-Margin-DeepVariant pipeline using recommended ONT parameters.

### Phasing of SNV and SVs

LongPhase was employed to phase the heterozygous SNVs and SV calls across the long reads. Longphase phase with ont settings was employed using the DeepVariant SNP/small indel calls along with the Sniffles SV calls. Phasing was then checked against parental haplotypes.

### ONT methylation calling and reference comparison

Nanopolish(Simpson et al. 2017) was used to call methylation profiles from the raw fast5 signal data. The fastq files were indexed to the corresponding fast5 signal data. Then, Nanopolish called methylation in CpG context from the data using the altered signal detected from incorporation of cytosine compared to 5mC. The initial output of detection per read was aggregated by genomic position using alignments to the reference genome; only reads with log likelihood >=2 were considered evidence of methylation. PycoMeth(Leger 2020) was used to find differentially methylated 1kb windows in proximity to genes and transcription start sites for affected vs unaffected individuals. Differentially methylated sites were compared to regulatory signature regions from ENCODE. NanoMethPhase was used to phase the methylated regions across haplotypes using the Pepper-Margin-DeepVariant SNV calls and WhatsHap (Martin, Ebert, and Marschall 2023) phasing.

### Transcriptome sequencing

RNA was extracted from the FFPE blocks using Qiagen RNeasy FFPE kit with Xylene deparaffinization. We extracted normal adjacent colon, 2 adenomas and the tumor sample of EO-CRC proband #2. Briefly, samples were lysed with proteinase K for 15 minutes then incubated at 80°C for 15 minutes. DNase treatment was applied to remove genomic DNA contamination. RNA was purified using RNeasy MinElute spin columns. Illumina libraries were prepared using Kapa stranded mRNA kit with RiboErase and Globin depletion. Libraries were prepared using unique barcode adapters. Libraries were pooled and were sequenced on NextSeq 2000 P2 PE101 dual indexed run.

Reads were mapped with STAR aligner (Dobin et al. 2013) to hg38 allowing shorter mapped lengths to account for the fragmented samples. Reads were assigned to genomic features (gene/exon GenCode annotations) using featureCounts (Y. Liao, Smyth, and Shi 2014). Normalization for sequencing depth and RNA composition as well as differential expression was performed using DESeq2 (Love, Huber, and Anders 2014).

## Supporting information

Supplemental Files List

## Data Access

Sequence data including raw signal fast5, aligned reads, and variant/methylation calls for the Oxford Nanopore PromethION and aligned reads and expression counts the Illumina RNA-seq will be accessible under a dbGap study (accession in progress). Institutional Review Board approval has been obtained from MSK by ZK Stadler.

## Acknowledgements

We thank the CSHL Cancer Center DNA Sequencing Shared Resource for providing the Oxford Nanopore and Illumina sequence data. The Sequencing Shared Resource is supported by the Cold Spring Harbor Cancer Center Grant (5P30CA045508). S.G. was supported by the National Institutes of Health (5R50CA243890). W.R.M is the Davis Family Professor of Human Genetics at CSHL. WRM would further like to acknowledge funding support from the CSHL/Northwell Health Affiliation for purchase of the ONT PromethION sequencer used in this study and the NIH 5P30CA045508 Cancer Center support grant. This work was in part funded by the Precision, Interception and Prevention Program (PIP) (Z.K.S); the Robert and Kate Niehaus Center for Inherited Cancer Genomics (Z.K.S. and K.O.) and the National Institutes of Health, National Cancer Institute Cancer Center Support Grants P30 CA008748 (all MSK authors) at Memorial Sloan-Kettering.

