## Supplemental Materials for "Exploring the genetic and epigenetic underpinnings of early-onset cancers: Variant prioritization for long read whole genome sequencing from family cancer pedigrees"

**Supplemental Materials Table of Contents**


Supplementary tables 1-8 and figures 1 and 2:

Sequencing coverage tables and histograms

Alignment summary tables

Structural variant summary

AnnotSV output tables

See files:
EOCRC_Trio1_proband_denovo.SV.annotated.tsv
EOCRC_Trio1_proband_hom.SV.annotated.tsv
EOCRC_Trio2_proband_denovo.SV.annotated.tsv
EOCRC_Trio2_proband_hom.SV.annotated.tsv
TGCT_Quad_BothSibsHet.SV.annotated.tsv
TGCT_Quad_BothSibsHom.SV.annotated

PycoMeth output tables

See files:
EOCRC_Trio1_pycoMeth_summary_intervals.tsv
EOCRC_Trio2_pycoMeth_summary_intervals.tsv
TGCT_Quad_pycoMeth_summary_intervals.tsv

Supplemental Figure 3

Supplemental Figure 4

*Supplemental Table1.*  *Sequencing quality metrics for Colon trio #1.*

**Colon Trio #1**
**Sequencing Stats**

| Sample | Total Q7 bases | Total Bases in reads >Q7/>30kb | Estimated coverage in reads >Q7/ >30kb | N50 readlength |
|---|---|---|---|---|
| C55A.2 (son) | 89,354,976,995 | 18,576,419,486 | 6X | 17795 |
| C55B.2 (father) | 73,079,756,690 | 32,783,695,190 | 10X | 27172 |
| C55C.2 (mother) | 105,208,921,400 | 33,670,870,780 | 10X | 22454 |

*Supplemental Figure 1.*  *Read-length histograms for ONT long read sequencing for Trio #1*

### Colon Trio #1 Sequencing Plots

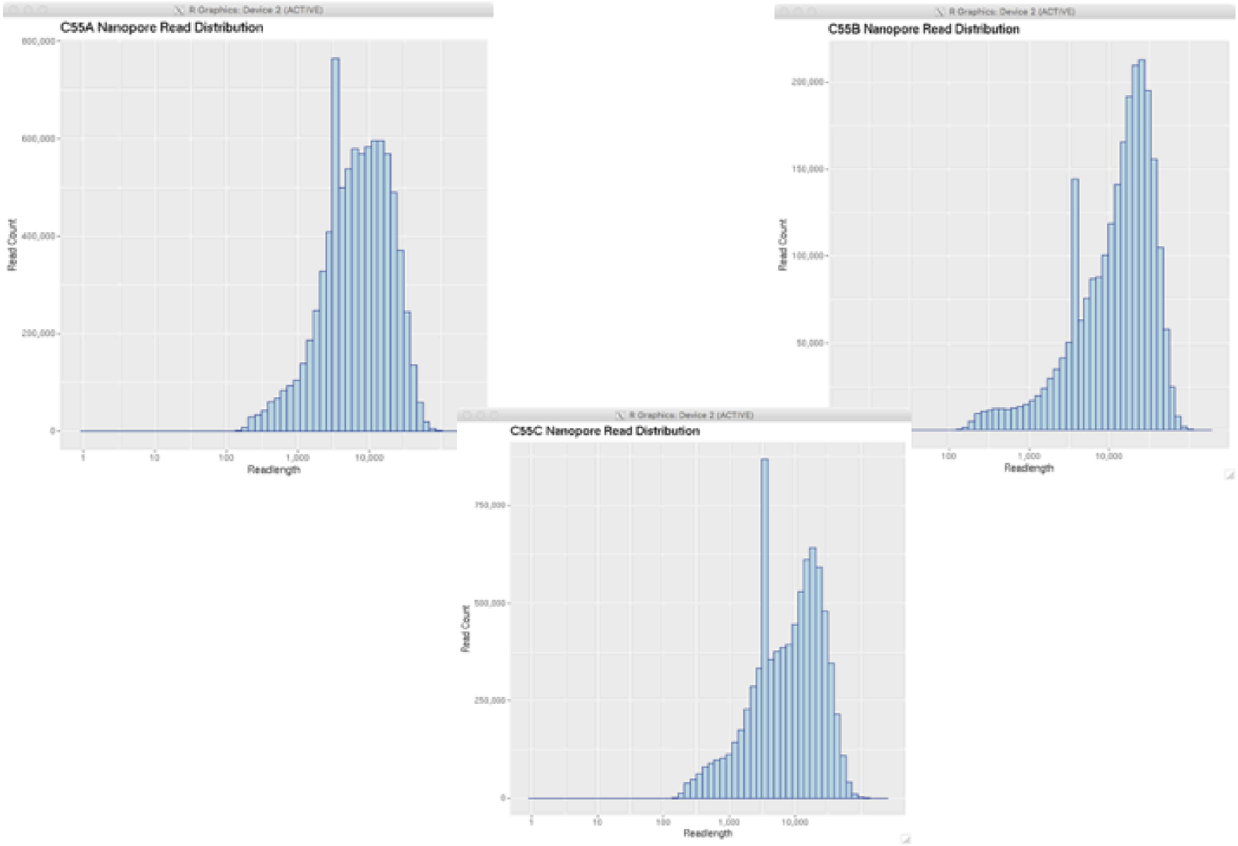


<u>*Supplemental Table 2.*</u> *Alignment metrics for colon trio 1.*

**Colon Family 1 (27797)**
**Alignment Stats**

| Sample | # Reads Mapped | % of Reads Mapped | Average Coverage |
|---|---|---|---|
| C55A.2 (son) | 7674073 | 90% | 25.0617 |
| C55B.2 (father) | 3719487 | 89% | 20.5696 |
| C55C.2 (mother) | 4748041 | 89% | 29.3589 |

*Supplementary Table 3.* *SV summary for colon trio #1.*

#### Colon Family(14-016, 09-068)
#### Variant Call Stats

| Sample | Total SV calls | Type |
|---|---|---|
| C55A.2<br>(son) | 24450 | 11791 DEL<br>1 DEL/INV<br>1613 DUP<br>21 DUP/INS<br>10529 INS<br>177 INV<br>342 TRA |
| C55B.2<br>(father) | 21001 | 9965 DEL<br>1 DEL/INV<br>1335 DUP<br>30 DUP/INS<br>9294 INS<br>145 INV<br>1 INVDUP<br>238 TRA |
| C55C.2<br>(mother) | 26448 | 12851 DEL<br>1 DEL/INV<br>1675 DUP<br>42 DUP/INS<br>11321 INS<br>201 INV<br>377 TRA |

*Supplemental Table 4.* *Alignment metrics for colon trio #2.*

**Colon Trio #2**
**Alignment Stats**

| Sample | # Reads Mapped | % of Reads Mapped | Average Coverage |
|---|---|---|---|
| SID103346 (Daughter) | 3935512 | 86% | 25.6797 |
| SID103347 (Mother) | 3329813 | 78.85% | 21.5947 |
| SID103348 (Father) | 4447138 | 87.81% | 30.4215 |

*Supplemental Table 5.* *SV summary for colon trio #2.*

### Colon Trio #2
### Structural Variant Stats

| Sample | Total SV calls | Type |
|---|---|---|
| SID103346 (Daughter) | 26620 | 11953 DEL<br>2273 DUP<br>11725 INS<br>243 INV<br>481 TRA |
| SID103347 (Mother) | 24386 | 11045 DEL<br>1962 DUP<br>10836 INS<br>206 INV<br>393 TRA |
| SID103348 (Father) | 27453 | 12314 DEL<br>2412 DUP<br>11939 INS<br>268 INV<br>567 TRA |

*Supplemental Table 6.* *Sequencing quality metrics for TGCT quad.*

#### Testicular Family 1 (14-016)
#### Sequencing Stats

| Sample | Total Q7 bases | Total Bases in reads >Q7/ >30kb | Estimated coverage in reads >Q7/ >30kb | N50 readlength |
|---|---|---|---|---|
| GCT-0267-1 Mother | 83,426,368,714 | 38,336,766,881 | 12X | 28404 |
| GCT-0267-11 Father | 121,020,159,105 | 44,263,180,995 | 14X | 24810 |
| GCT-0267-12 Son (aff) | 86,002,749,997 | 31,188,431,665 | 10X | 25087 |
| GCT-0267-21 Son (bilateral) | 123,788,146,309 | 27,511,340,879 | 9X | 21871 |

*Supplementary Figure 2.* *Read length histograms for ONT sequencing for TGCT quad.*

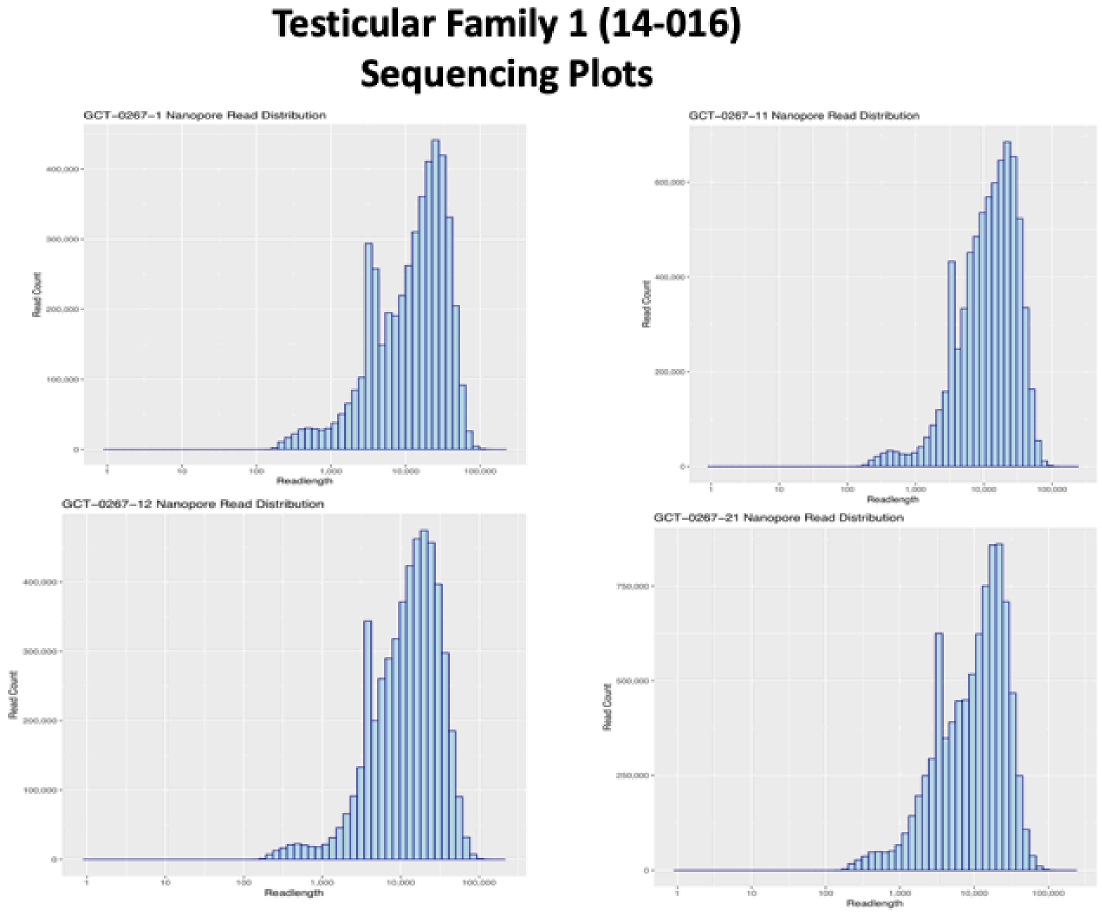


*Supplemental Table 7.* *Alignment metrics for TGCT quad.*

#### Testicular Family 1 (14-016)
#### Alignment Stats

| Sample | # Reads Mapped | % of Reads Mapped | Average Coverage |
|---|---|---|---|
| GCT-0267-11 Father | 6733410 | 91% | 34.0743 |
| GCT-0267-1 Mother | 4108296 | 87% | 23.6041 |
| GCT-0267-21 Son (bilateral) | 7985885 | 91% | 34.8088 |
| GCT-0267-12 Son | 4652843 | 91% | 24.1907 |

*Supplemental Table 8.* *SV summary for TGCT quad.*

#### Testicular Family 1 (14-016)
#### Variant Call Stats

| Sample | Total # Calls | Type |
|---|---|---|
| GCT-0267-1<br>Mother | 23353 | 11243 DEL<br>1479 DUP<br>26 DUP/INS<br>10167 INS<br>160 INV |
| GCT-0267-11<br>Father | 27647 | 13237 DEL<br>1 DEL/INV<br>1794 DUP<br>45 DUP/INS<br>11742 INS<br>281 INV |
| GCT-0267-12<br>Son | 23855 | 11851 DEL<br>1 DEL/INV<br>1413 DUP<br>21 DUP/INS<br>10097 INS<br>188 INV |
| GCT-0267-21<br>Son (bilateral) | 28755 | 14388 DEL<br>1 DEL/INV<br>1815 DUP<br>39 DUP/INS<br>11720 INS<br>276 INV |

*Supplemental Figure 3. IGV screenshot of EO-CRC trio #1 aligned to CHM13. Aligned read panels show homozygous deletion in proband, heterozygous deletion in the unaffected parents. UCSC Genome Browser track shows a break in the alignment between CHM13 and hg38 in this region.*

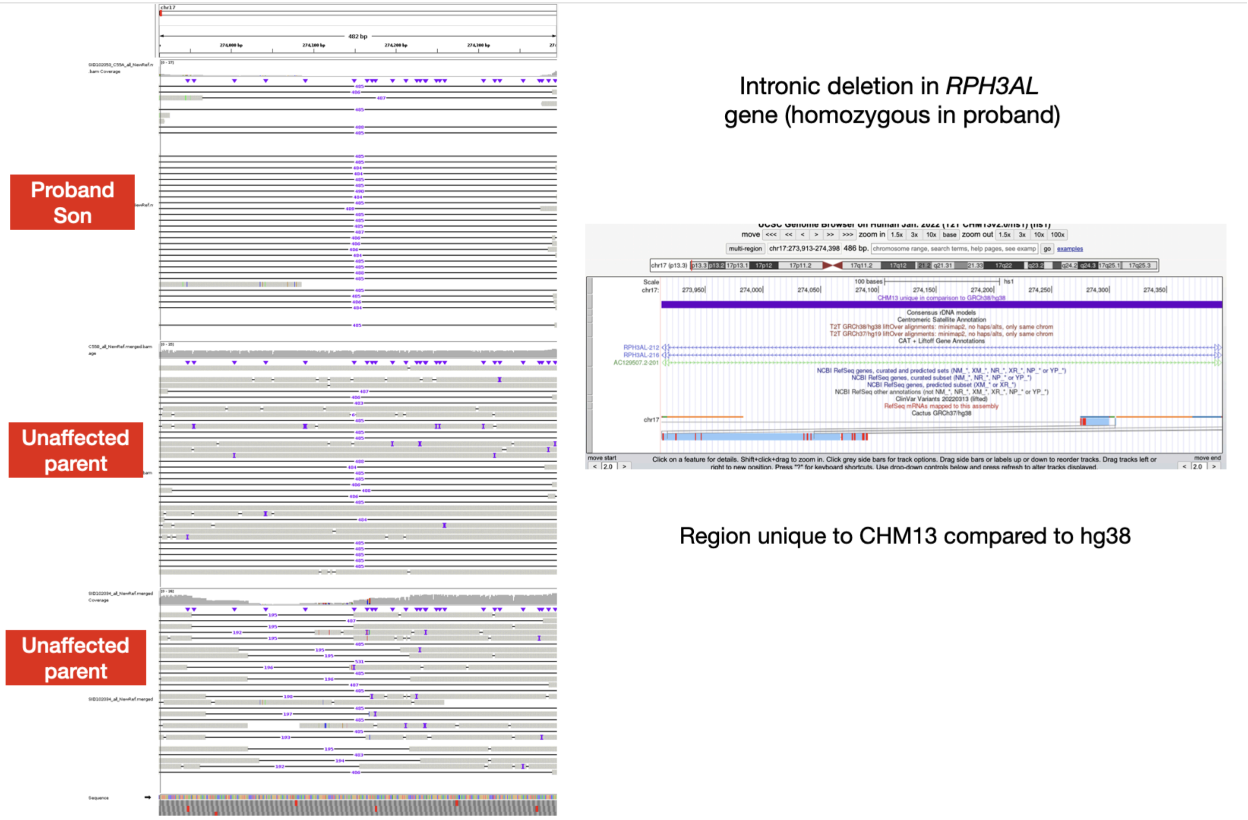


*Supplemental Figure 4. Expression counts in selected genes harboring SVs in the proband compared to healthy parents. Normalized counts (denoted as circles, 3 replicates per sample) are shown for FFPE extractions of 3 groups - normal adjacent colon (normal), two different adenomatous polyps (polyp), and colon tumor (tumor).*

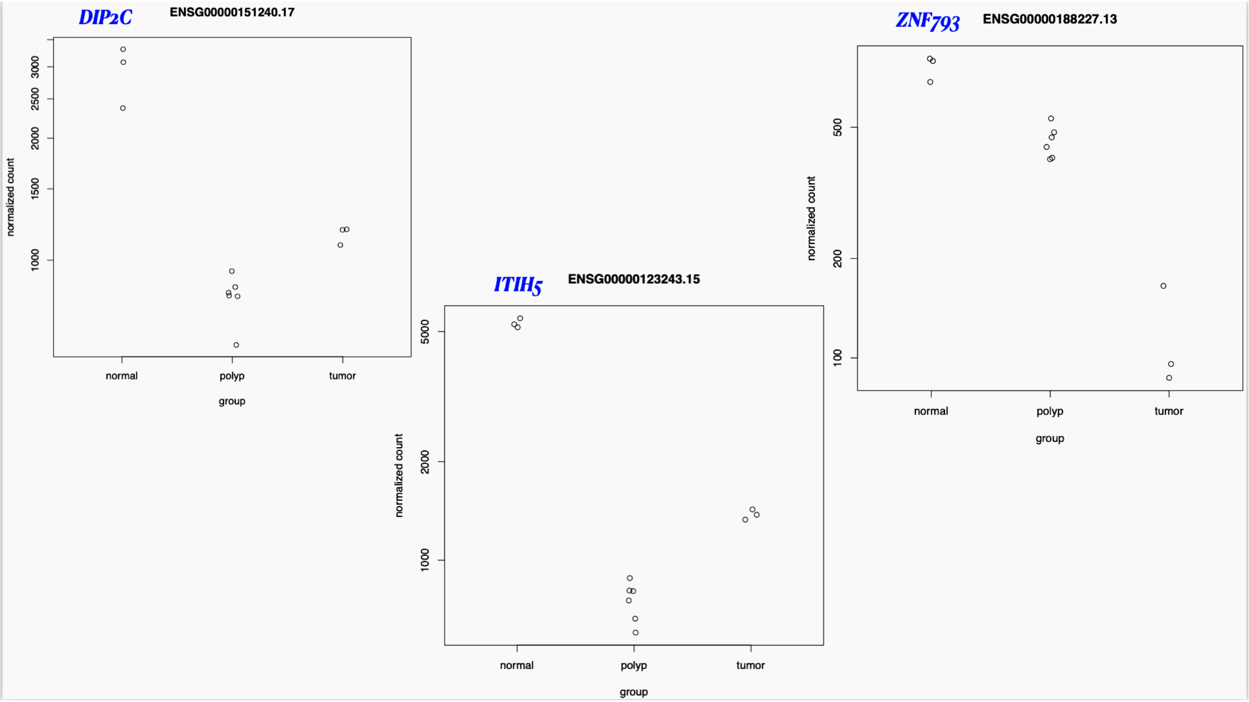
DIP2C
ENSG00000151240.17
normalized count
1000
1500
2000
2500
3000
normal
polyp
tumor
group
ITIH5
ENSG00000123243.15
normalized count
1000
2000
5000
normal
polyp
tumor
group
ZNF793
ENSG00000188227.13
normalized count
100
200
500
normal
polyp
tumor
group
